## Supplementary Information for "Oculomotor freezing reflects tactile temporal expectation and aids tactile perception"

Badde et al.

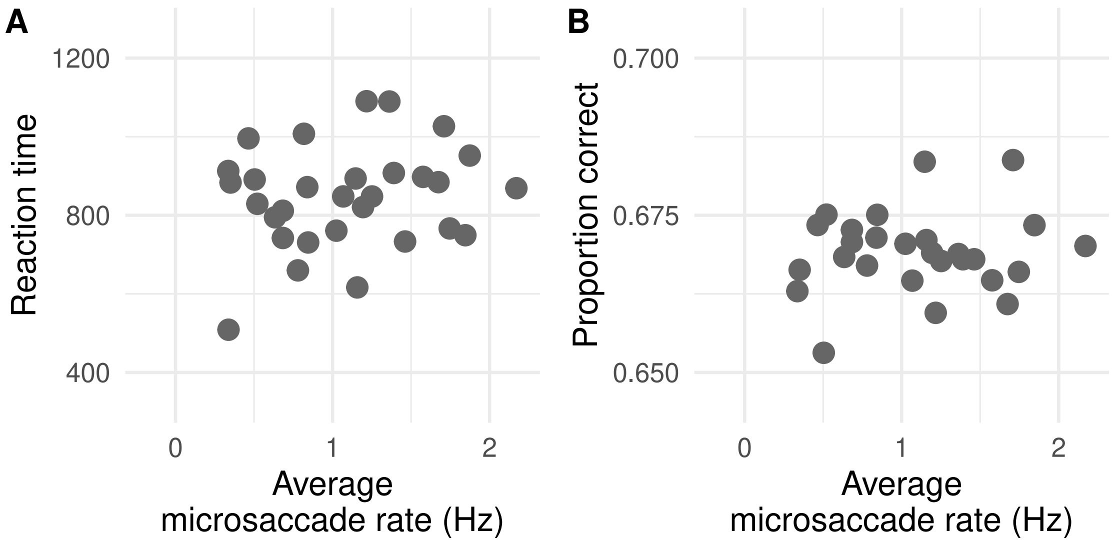


Supplementary Figure 1. Average task performance and microsaccade rates. Participants' average reaction time (A) and proportion of correct responses (B) as a function of their average microsaccade rate during the 1000-ms long interval before target onset. Please note, data of three participants with lower proportions of correct values (0.53, 0.55, 0.60) are not included in (B) to avoid leverage effects. Otherwise, the figures are based on the full dataset (N = 30 participants, 100 repetitions per all 10 conditions and each participant). Source data are provided as a Source Data file

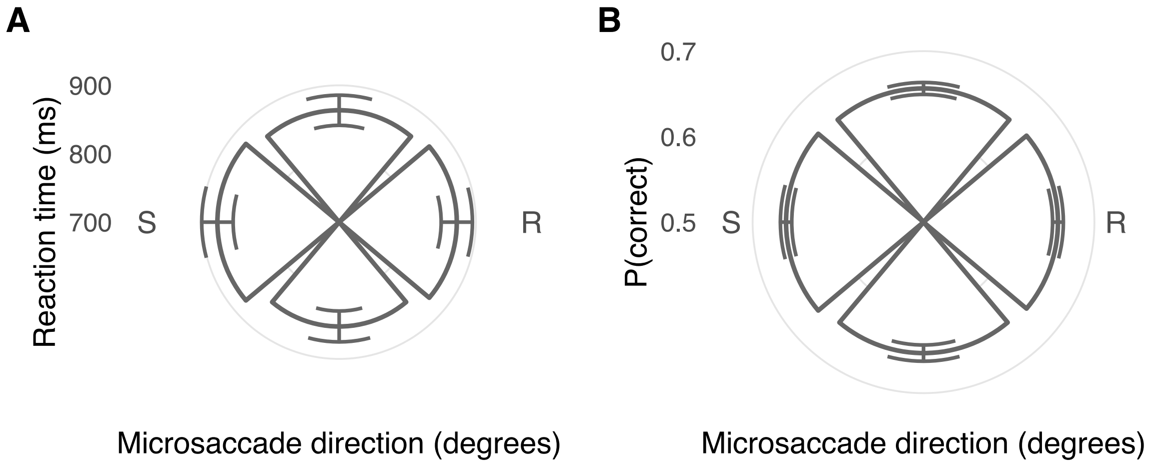


Supplementary Figure 2. Average task performance and microsaccade direction. Group mean reaction time (A) and proportion of correct responses (B) in trials in which a microsaccade occurred during the cue-target-interval as a function of microsaccade direction (S stimulated hand; R response hand). Error bars show standard errors of the mean. All statistics are based on the full dataset (N = 30 participants, 100 repetitions per condition and participant) and source data are provided as a Source Data file.

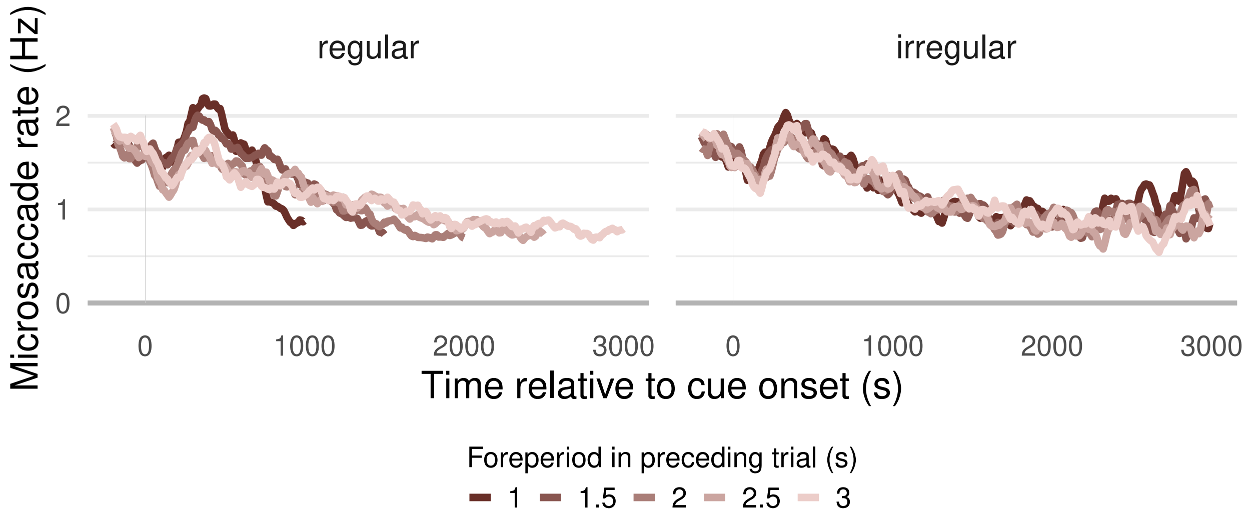


Supplementary Figure 3. Microsaccade rates and sequential foreperiod effects. Group mean microsaccade rates as a function of trial time relative to the onset of the tactile cue separately for each predictability condition (panels) and foreperiod in the preceding trial (red shades). Timelines were cut off at the onset of the tactile target stimulus to allow averaging across foreperiods in the current trial in the irregular condition. In the regular condition, the foreperiod in the preceding trial is by definition identical to the foreperiod in the current trial. All shown group mean values are based on the full dataset (N = 30 participants, 100 repetitions per condition and participant) and source data are provided as a Source Data file.

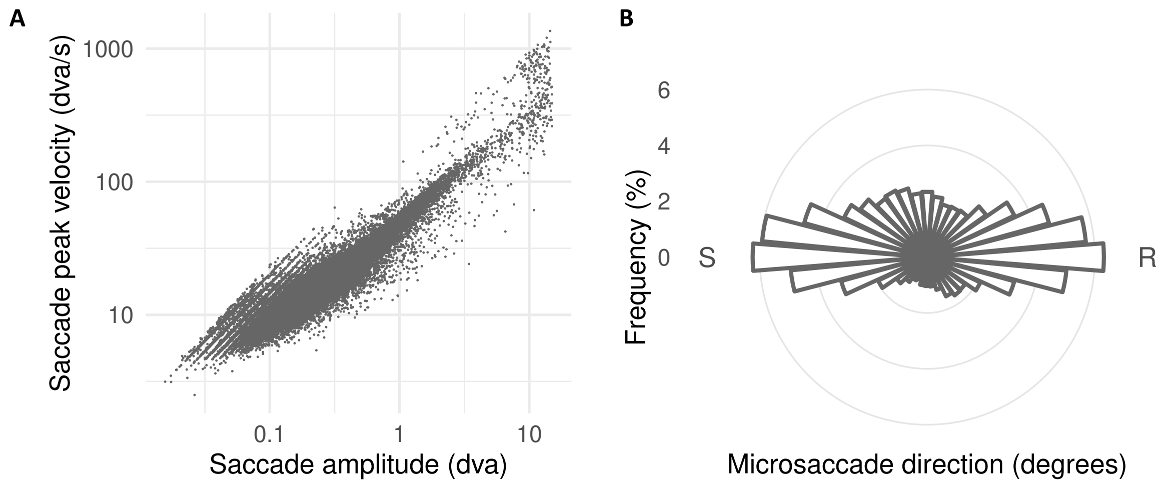


Supplementary Figure 4. (A) Saccade amplitude and peak velocity. Microsaccades and saccades followed the main sequence. For results reported in the manuscript, only microsaccades, i.e., saccades with an amplitude below 1 dva were analyzed. (B) Microsaccade direction. Microsaccade directions during the time interval between cue and target stimulus relative to the location of the stimulated (S) or the response (R) hand. Both panels are based on the full dataset (N = 30 participants, 100 repetitions per condition and participant), (A) shows parameters of individual saccades, (B) shows group mean frequencies. Source data are provided as a Source Data file

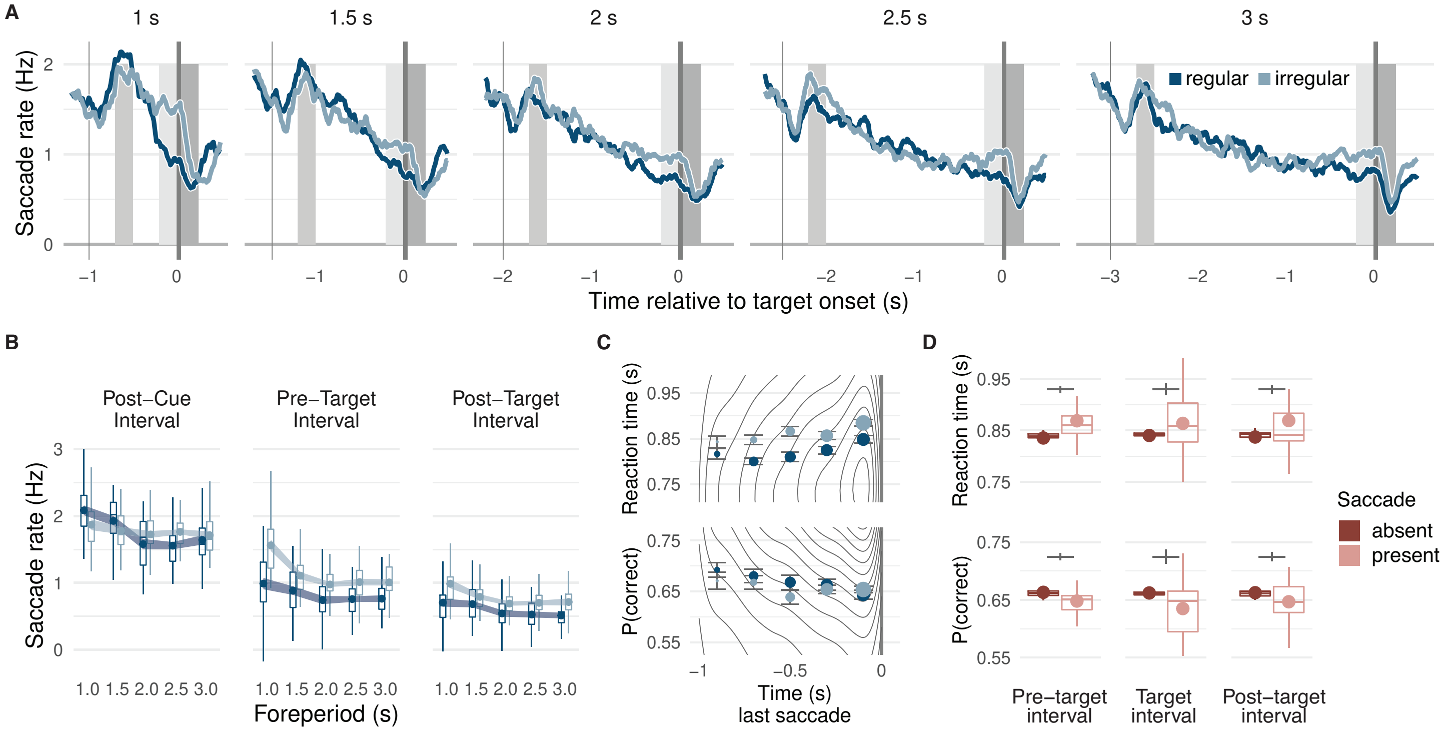


Supplementary Figure 5. Saccade rates, temporal predictability, and task performance. (A) Group mean saccade rates as a function of trial time relative to the onset of the tactile target stimulus separately for each predictability condition (dark blue, regular; light blue, irregular) and foreperiod (panels). Shaded vertical bars indicate the cue and target stimulus (blackish grey), shaded rectangles the post-cue (medium grey), pre-target (light grey), and post-target (dark grey) intervals. (B) Average saccade rates in post-cue, pre-target, and post-target intervals. Boxplots show the distribution of participant-level mean values per condition adjusted by their overall interval mean (center line, median; box limits, upper and lower quartiles; whiskers, minimum and maximum limited by 1.5x interquartile range). Circular markers show group-level mean values; the width of the ribbon matches the predictability-condition-adjusted standard error. (C) Task performance by saccade latencies. Grey lines indicate the 2d frequency distribution of single trial reaction times (upper panel) and correct responses (lower panel) as a function of the latency of the last saccade before the target stimulus. Circular markers show group mean reaction times in regular (dark blue) and irregular (light blue) conditions for 200-ms long time bins. Marker size represents the percentage of trials per bin. Error bars indicate standard errors corrected for between-participant variability. (D) Task performance and saccades. Reaction times (upper panel) and proportion correct (lower panel) split by the presence of saccades in the pre-target, target, and post-target interval (dark red, absent; light red, present). Boxplots (defined as in B) indicate the distribution of participant-level mean values per condition adjusted for their overall mean. Please note that trials without a microsaccade in the respective interval were more frequent than those with a microsaccade resulting in a narrower distribution. Circular markers show group-level mean values. Vertical grey lines indicate the standard error of the difference between conditions. All statistics are based on the full dataset (N = 30 participants, 100 repetitions per condition and participant) and source data are provided as a Source Data file.

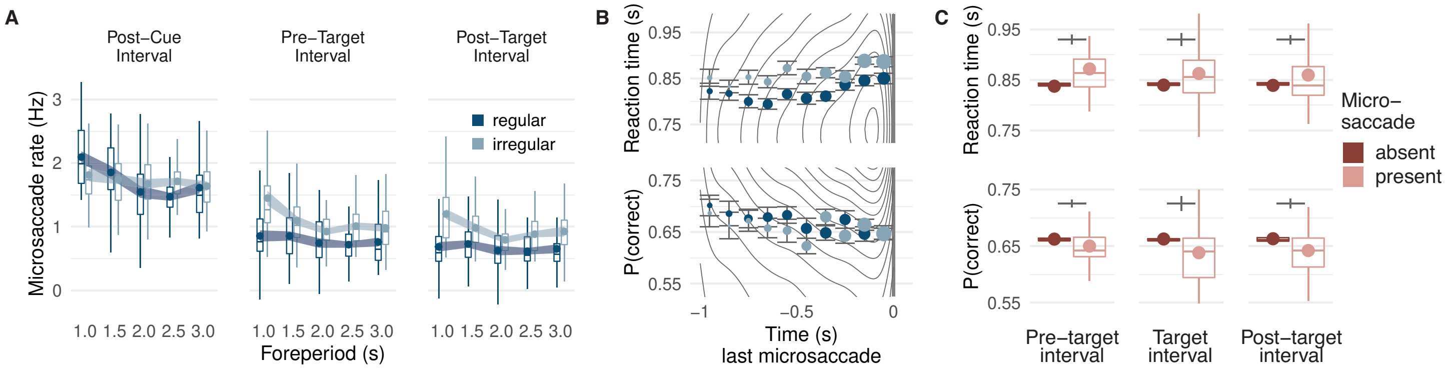


Supplementary Figure 6. Microsaccade rates within shorter intervals of interest, temporal predictability, and task performance. (A) Microsaccade rates in a comparison interval 400-500 ms after the onset of the tactile cue (left panel), in the 100 ms interval before the onset of the tactile target stimulus (center panel), and in a post-target interval 0-100 ms after the offset of the tactile stimulus (right panel), separately for each predictability condition (regular: dark blue; irregular: light blue) and foreperiod (x-axis). Boxplots indicate the distribution of participant-level mean values per condition adjusted by their overall interval mean (center line, median; box limits, upper and lower quartiles; whiskers, minimum and maximum limited by 1.5x interquartile range). Circular markers show group-level mean values; the width of the ribbon matches the predictability-condition-adjusted standard error which indicates the degree of inter-subject variation in the difference between regular and irregular conditions and therefore whether there is a significant effect of predictability condition on microsaccade rates. (B) Task performance by microsaccade latencies. Grey lines indicate the 2d frequency distribution of single trial reaction times (upper panel) and correct responses (lower panel) as a function of the latency of the last microsaccade before the target stimulus. Circular markers show group mean reaction times in regular and irregular conditions for 100-ms long time bins. Marker size represents the percentage of trials per bin. Error bars indicate standard errors corrected for between-participant variability. (C) Task performance and microsaccades. Group mean reaction times (upper panel) and proportion correct (lower panel) split by the presence of a microsaccade in the pre-target, target, or post-target interval. Boxplots (defined as in A) indicate the distribution of participant-level mean values per condition adjusted for their overall mean. Please note that trials without a microsaccade in the respective interval were more frequent than those with a microsaccade resulting in a narrower distribution. Circular markers show group-level mean values. Vertical grey lines indicate the standard error of the difference between conditions. All statistics are based on the full dataset (N = 30 participants, 100 repetitions per condition and participant) and source data are provided as a Source Data file.
